# Rare variation in malaria parasites biases population-genetic inference

**DOI:** 10.64898/2026.01.09.698000

**Authors:** Amy Goldberg

## Abstract

Understanding how pathogens evolve is fundamental to disease control and is a basic question in evolutionary biology, yet pathogens with complex life cycles violate assumptions of classic evolutionary models. Genetic analyses of the malaria parasite *Plasmodium falciparum* have shown multiple surprising patterns. For example, empirical analyses produce effective population size estimates that vary by orders of magnitude depending on the method and an excess of genes with elevated nonsynonymous variation (measured as *π*_*N*_ /*π*_*S*_). Here, reanalyzing genomic data from 18 worldwide populations, I show that these observations directly follow from distributions of genetic variation enriched for rare variants. Multiple potential mechanisms may increase the proportion of rare variants, including host expansions, lifecycle dynamics, population structure, or selection. The observed genealogies are more consistent with a multiple-merger coalescent than a Kingman coalescent, causing common summary statistics to be biased in predictable directions. In particular, the abundance of rare variants interacts with the mathematical properties of ratio statistics to systematically inflate gene-level *π*_*N*_ /*π*_*S*_ estimates even in the absence of selection. Notably, filtering this rare variation reveals previously masked candidates for selection, including well-characterized antigens such as merozoite surface proteins. This framework provides a foundation for interpreting genomic data in pathogens with high variance in reproductive success.

**Significance Statement:** Pathogen genomics increasingly guides public health decisions, yet organisms with complex life cycles can produce genealogies that violate assumptions of standard population genetic methods. Here I show that the malaria parasite *Plasmodium falciparum* has an excess of rare variation, perhaps produced from overdispersed distributions of reproductive success. These genealogies generate systematic biases in commonly reported statistics arising from the interaction between rare variation and the mathematical properties of the statistics themselves, not from biological processes often invoked to explain anomalous patterns. Correcting for this bias reveals signatures of selection in well-characterized vaccine candidates that were previously masked, demonstrating that appropriate null models are essential for identifying targets of intervention and inferring evolutionary history in pathogens and other systems with reproductive skew.

## 1 Introduction

Parasite genomic information is increasingly used to guide public health decisions and clinical treatment, based on recent advances in phylogenetic and population genetic methods (1). However, pathogens with complex life cycles, e.g. involving sexual reproduction and extreme demographic variation, have largely been left behind.

For example, tens of thousands of whole genomes are now available for the malaria parasite *Plasmodium falciparum* (2; 3), and recent theoretical studies have demonstrated that the malaria life cycle intensifies genetic drift and the efficacy of selection (4; 5) and have made made major progress building frameworks for predictive genetic epidemiology to guide public health decisions based on genetic data (6; 7; 8). Yet multiple empirical observations from these data remain unexplained or potentially contradictory, leading to calls for progress on modeling genetic variation of the parasites (9; 10). First, estimates of effective population size vary dramatically from 10^2^ to 10^7^ depending on methodology (11; 12; 13). Second, approximately 20-30% of *P. falciparum* genes exhibit π_N_ /π_S_ > 1, nearly ten times the proportion observed in *Drosophila melanogaster* or humans where this fraction remains below 5% (4; 14; 15). This pattern has been interpreted as relaxed constraint leading to accumulation of slightly deleterious mutations (potentially lifecycle-stage specific) or as increased purifying selection on synonymous mutations (4; 14; 15; 10).

Here, I propose that these perhaps puzzling patterns may arise from the genealogical structure of *P. falciparum*, which follows patterns of a multiple-merger coalescent, in which multiple lineages trace back to one common ancestor in a single generation. Then, I consider the implications of this model for interpretation of summaries of genetic variation.

Within a single transmission cycle, parasites undergo massive expansions and bottlenecks providing opportunities to skew reproductive success and increasing the chance of multiple merger coalescence (10; 16; 15). At the population level, parasites experience variation in transmission success among hosts, with some infections producing many onward transmissions while most produce few or none. This between-host variance in reproductive success shapes population-level genealogies and has direct consequences for the site frequency spectrum and summary statistics derived from it.

When offspring distributions are highly skewed, the Kingman coalescent, which assumes only pairwise mergers occur, is a poor fit. Instead, these genealogies follow multiple-merger coalescence patterns, where many lineages find a common ancestor in a single event. Multiple frameworks for a multiple-merger coalescent have been proposed (17; 18; 19; 20; 21; 22; 23), and they are increasingly appreciated as representative of the genealogy of many empirical populations (23; 24; 25). One well-studied version is the Beta coalescent, parameterized by *α* ∈ (1, 2], where *α* = 2 recovers the Kingman coalescent and smaller *α* indicates increasingly skewed offspring distributions. Intuitively, small *α* means a few individuals each generation leave disproportionately many descendants, causing bursts of coalescence when tracing ancestry backward. The observable consequence is an excess of rare variants: mutations arising on rapidly expanding branches that have not yet drifted to higher frequency.

## 2 Empirical fit of a Beta coalescent

Site frequency spectrum (SFS) from each of 18 MalariaGEN *Pf8* populations showed an excess of rare variants compared to a standard Kingman coalescent (Figs. 1,S1,S2, Table S1). Under the Beta coalescent with 1 < *α* < 2, rare variants are enriched relative to Kingman expectations, with the degree of enrichment increasing as *α* decreases. Therefore, I fit Beta(2 − *α, α*) coalescent models to the SFS of each population (Fig. 1). I also compared the fit of a Kingman-like coalescent with exponential population size growth, as an alternative model that is also expected to result in an excess of rare variants (Table S2).

**Figure 1.**
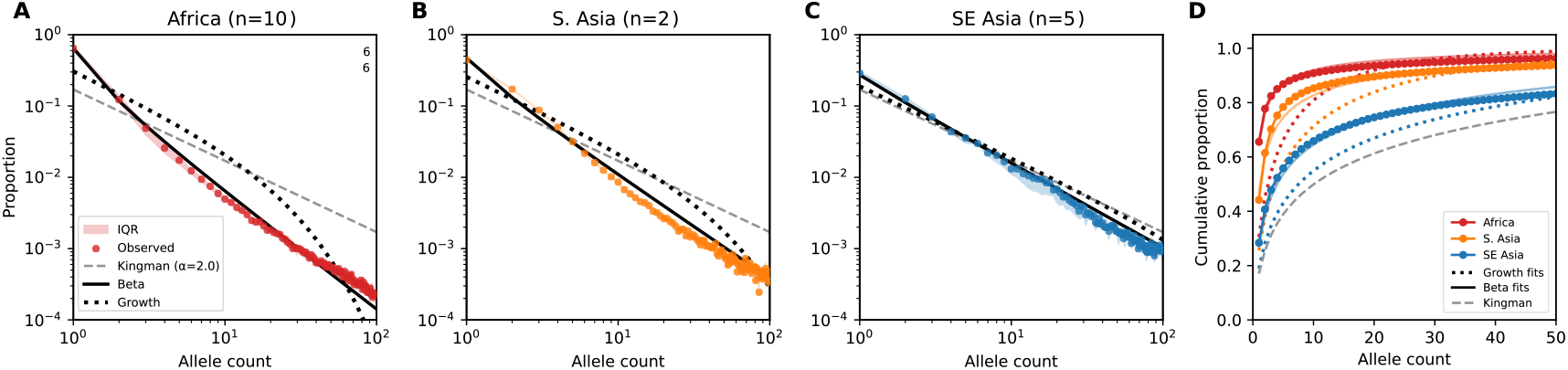
Empirical genomic data from *P. falciparum* fit a Beta coalescent model. **(A–C)** Observed SFS (colored points) on a log-log scale compared to fitted Beta coalescent (black line), Kingman coalescent with exponential growth (black short-dashed line), and Kingman coalescent expectations (*α* = 2.0; dashed gray line) for African, South Asian, and Southeast Asian populations, respectively, from MalariaGEN *Pf8*. Color shaded regions in (A, C) indicate interquartile range across populations. Regional *α* values are maximum likelihood estimates averaged across constituent populations (n = number of populations). **(D)** Cumulative distributions of the empirical SFS for mean-by region compared to Beta coalescent fit per-region, Kingman coalescent with exponential growth fit per-region, and the Kingman coalescent. All populations reject the Kingman coalescent (likelihood ratio tests, *p* < 0.001)

I estimated *α* by maximum likelihood using the Poisson random field approximation (26), with expected SFS proportions under the Beta(2 − *α, α*) coalescent from Birkner et al. (2013) (21) (Materials & Methods). Figure 1 shows the observed site frequency spectrum for each region compared to Kingman and Beta coalescent expectations. Across all regions, the observed SFS shows a pronounced excess of rare variants relative to Kingman predictions (dashed gray), with the Beta coalescent providing substantially better fit. The excess is most extreme in African populations (Fig. 1A), where singletons comprise 61–72% of segregating sites compared to < 20% expected under the Kingman coalescent for *n* = 200. African *P. falciparum* populations showed significantly stronger multiple-merger signatures 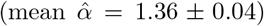 than South Asian 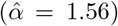 and Southeast Asian populations 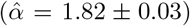, with all populations deviating significantly from the standard Kingman coalescent (all likelihood ratio tests *p* < 0.001; Table S2).

In some low-transmission populations, the observed SFS is consistent with either weak Beta coalescent (*α* ≈ 1.8) or moderate exponential growth. These populations may experience population-level expansion (e.g. through changes in case numbers or expansion of drug resistant lineages) or may simply exhibit lower multiple-merger dynamics due to lower infection rates (e.g. lower multiplicity of infection (MOI) reducing co-transmission bottleneck intensity). For Laos and PNG in particular, exponential growth and Beta coalescence produce similar SFS shapes as *α* approaches 2, making them difficult to distinguish.

African populations exhibit both higher diversity and stronger multiple-merger signatures than Southeast Asian populations. Under the Beta coalescent, smaller *α* accelerates coalescence; therefore, the occurrence of higher diversity in Africa despite smaller *α* requires a substantially larger underlying metapopulation effective size. This is consistent with Africa’s higher transmission intensity, greater number of concurrent infections, and elevated multiplicity of infection.

The stronger fit of the Beta coalescent (*α* ≪ 2) within endemic regions suggests that *P. falciparum* evolution there is governed by high variance in reproductive success, where a small minority of infections contribute disproportionately to the next generation. The geographic gradient in 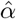 provides insight into multiple potentially co-occurring mechanisms. For example, high vector density may allow a high-gametocyte host to infect many of mosquitoes per day. Further, higher average MOI both increases transmission events that carry multiple related lineages and amplifies the impact of stochastic loss through transmission bottlenecks. In Southeast Asia, reduced MOI and fewer vectors may decrease this variance, causing lineages to coalesce in a more pairwise fashion closer to Kingman coalescence.

## 3 Bias in summaries of neutral variation

Given the fit of the Beta coalescent model to empirical *P. falciparum* data, I next consider the implications for inference from population-genetic summary statistics. Descriptive summary statistics are commonly used to interpret empirical malaria genetics data. In particular, a classic signature of Beta coalescence is an excess of rare variants compared to a Kingman coalescent. The skewed SFS has implications for interpretation of summary statistics based on the SFS, and for inference of population history or selection from SFS.

### 3.1 Tajima’s *D*

Tajima’s *D* compares two estimators of genetic diversity, *θ*_*π*_ (mean pairwise differences) and *θ*_*W*_ (based on segregating sites), and is sensitive to recent demographic history and selection. In population samples, negative Tajima’s *D* values indicate an excess of rare variants, typically interpreted as a signature of rapid population expansion or positive selection, while positive values may indicate balancing selection or population structure. Under a constant-size Wright-Fisher model, Tajima’s *D* is expected to be centered on zero. However, empirical malaria genetic studies consistently show genome-wide Tajima’s *D* values that are highly negative (14; 15; 27; 28; 29; 30; 31; 32; 33).

In the *Pf8* dataset, Tajima’s *D* values are strongly negative across all populations, consistent with the rare variant excess predicted by Beta coalescent dynamics (Fig. 2, Table S3). Genomewide *D* ranges from −1.5 to −2.5 across populations, values that would traditionally be interpreted as strong population expansion or positive selection, but instead can be produced by the multiple merger genealogical structure without needing to invoke selection or host-population expansions.

**Figure 2.**
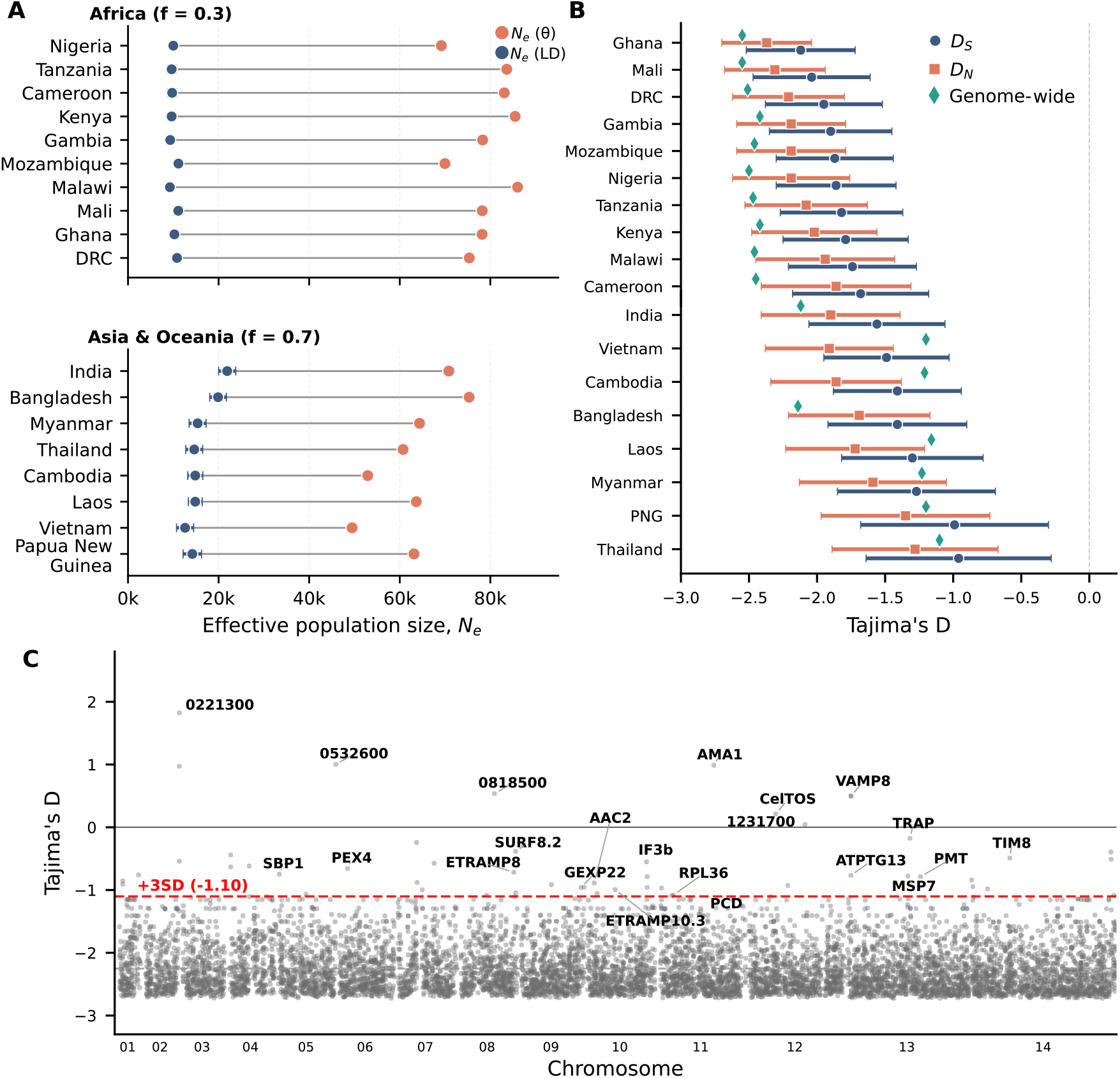
Summary statistics biased rare-variant excess in empirical *P. falciparum* data. **(A)** Difference in effective population size, *N*_*e*_, estimated from nucleotide diversity versus linkage disequillibrium (LD). Across all populations, nucleotide-diversity–based *N*_*e*_ estimates are consistently several-fold larger than LD–based estimates. Values in Table S4. **(B)** Tajima’s *D* by population. Genome-wide Tajima’s *D* is strongly negative in all populations (green diamonds). The mean of per-gene Tajima’s *D* is consistently negative, though the values are less extreme than genome-wide *D* in African populations, whereas in some low-diversity Asian populations, pergene *D* is more negative than the genome-wide estimate. *D*_*N*_ and *D*_*S*_ are calculated using only nonsynonymous and synonymous variants, respectively. Values in Table S3. **(C)** In a representative high-transmission population (Ghana), the distribution of per-gene Tajima’s *D* is shifted far below zero, yet positive outliers still recover known antigenic and vaccine-candidate genes using 3 standard deviations (*D >* −1.1) as a cutoff.

Genome-wide *D* and the mean of per-gene *D* estimates differ (Fig. 2B). These two quantities weigh genes differently: genome-wide *D* pools all segregating sites before computing the statistic, so genes with more polymorphisms contribute proportionally more information. The mean of pergene *D* weights all genes equally, regardless of how many segregating sites each contains. Genes with few segregating sites yield unstable *D* estimates with high variance, yet contribute equally to the per-gene average. In African populations, genome-wide *D* is typically more negative than per-gene means; the pooled estimate captures the strong rare-variant signal cleanly, while pergene averaging is diluted by noisy estimates from low-diversity genes. In Asian populations with lower overall diversity, this pattern can reverse; many genes contain only a handful of singletons, producing extreme negative *D* values that pull the per-gene mean below the genome-wide estimate. This sensitivity to aggregation method under Beta coalescent genealogies has broader implications for summary statistics beyond Tajima’s *D* (see below for discussion for *π*_*N*_ /*π*_*S*_).

Despite the strongly negative genome-wide values, per-gene Tajima’s *D* scans can identify candidates for balancing selection as positive outliers. In Ghana, the distribution is sufficiently shifted that even a +3 standard deviation threshold remains negative (*D* = −1.10; Fig. 2C). Genes exceeding this threshold include well-characterized vaccine candidates and surface antigens: AMA1 (*D* = 0.99), CelTOS (*D* = 0.21), TRAP (*D* = −0.18), SBP1 (*D* = −0.75), and MSP7 (*D* = −0.78).

These genes encode proteins under strong immune pressure, where frequency-dependent selection is expected to maintain antigenic polymorphism. Their appearance as outliers, despite falling within the range that would be considered neutral or negative in other organisms, demonstrates that selection scans remain informative despite SFS skew under an appropriate null expectation.

### 3.2 Estimators of *N*_*e*_

Next, I consider multiple estimators for effective population size, *N*_*e*_. I calculated *N*_*e*_ using both nucleotide diversity-based and linkage disequilibrium (LD)-based approaches. Under the standard neutral model, expected nucleotide diversity equals the population-scaled mutation rate: 𝔼 [*π*] = *θ* = 2*N*_*e*_*µ* for haploid organisms. Diversity-based estimates using this formula ranged from 49,474 to 86,019 (median = 73,078) across populations. LD-based estimates produced substantially smaller values, 9,247 to 21,898 (median = 11,119). Fig. 2A and Table S4 show that the ratio of diversity-based to LD-based estimates was consistently 3–9-fold across populations (median 6.6).

This divergence is expected when the underlying genealogy follows a multiple-merger process. Under a Kingman coalescent dynamics diversity scales linearly with effective population size. Under a Beta coalescent with *α* < 2, however, this relationship becomes sublinear, with 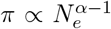 (21). Multiple-merger events cause coalescence times to grow less than proportionally with population size, so observed diversity is lower than Kingman predictions for a given *N*_*e*_. Conversely, applying Kingman-based formulas to diversity generated under a Beta coalescent produces higher *N*_*e*_ estimates. With *α* ≈ 1.36 in African populations, diversity scales as 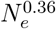 rather than linearly; an effective size of ∼10, 000 would produce diversity consistent with *N*_*e*_ ≈ 60, 000–80,000 under Kingman assumptions, broadly matching the observed inflation.

The consistent divergence between these estimators is therefore expected when the underlying genealogy follows a multiple-merger process. The two estimators differ in which portion of the allele frequency spectrum they sample. LD-based estimators are driven by correlations among alleles at intermediate frequencies; rare variants contribute negligibly to *r*^2^ because singletons cannot generate measurable linkage disequilibrium. In contrast, nucleotide diversity integrates information across the full frequency spectrum, including the rare variants that distinguish Beta from Kingman coalescent dynamics. The rare-variant-enriched site frequency spectrum inflates diversity-based estimates when interpreted through Kingman formulas, while LD-based estimates remain closer to Kingman expectations because they effectively condition on common variation.

Notably, the ratio of diversity-based to LD-based *N*_*e*_ tends to be larger in African populations than in Southeast Asian populations, paralleling the stronger multiple-merger signatures (smaller 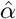) inferred from the SFS. This correlation provides additional support for the interpretation that genealogical structure, rather than demographic history alone, underlies the divergence between estimators. These results caution against interpreting any single *N*_*e*_ estimate as representing a true population parameter in systems with skewed offspring distributions. Each estimator captures information about different aspects of the underlying genealogical process, and their divergence may itself be diagnostic of multiple-merger dynamics.

## 4 Excess of rare variation combined with mathematical properties of ratio statistics inflate gene-level *π*_*N*_ /*π*_*S*_

The excess of rare variants, consistent with a Beta coalescent, has important statistical consequences for interpretation of common population-genetic summary statistics beyond the SFS. Here, I consider the impact on gene-level *π*_*N*_ /*π*_*S*_ estimates because of their historical interpretation in empirical and theoretical malaria genomic papers.

The observation that ∼20–30% of *P. falciparum* genes exhibit *π*_*N*_ /*π*_*S*_ *>* 1 was a striking departure from other organisms, where this fraction remains below 5% (4; 14; 15; 34). These patterns have been attributed to increased purifying selection on synonymous variation owing to the large within-host expansion size or to stage-specific relaxation of constraint from drift (4; 10). Here, I suggest that these patterns are largely explained by statistical properties of *π*_*N*_ /*π*_*S*_ as a ratio under the excess of rare variants of a Beta coalescent model, combined with low overall diversity for Southeast Asian populations.

### 4.1 Mathematical properties of ratio statistics

Calculating genome-wide values across the 18 populations from MalariaGEN *Pf* 8, *π*_*N*_ /*π*_*S*_ = 0.56 *±* 0.04. Yet, the mean of gene-level *π*_*N*_ /*π*_*S*_ ratios ranges from 1.5 to 8.6, 3- to 15-fold higher than genome-wide estimates (Fig. 3).

**Figure 3.**
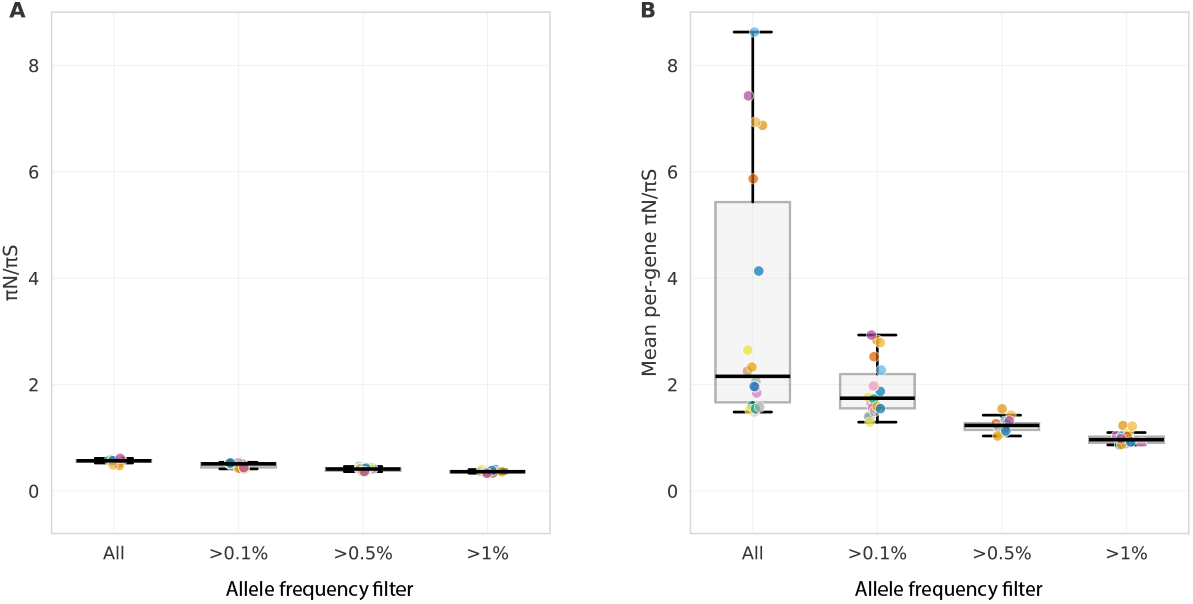
The mean of ratios is larger than the ratio of means for diversity in *Plasmodium*. **(A)** plots the genome-wide quantity 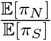, **(B)** plots the mean per-gene quantity 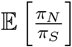. Different values of allele frequency filtering are considered along the x axis: all variants included, or only variants with a within-population frequency of the listed number. Following Jensen’s inequality, genome-wide values in panel A are always less than the values of the same country in panel B. Allele frequency filtering decreases this discrepancy. Values in Table S5.

The statistical properties of genome-wide quantities are distinct from per-gene comparisons. Following Jensen’s inequality, for a convex function *f*, 𝔼 [*f* (*X*)] ≥ *f* (𝔼 [*X*]), with equality only when *f* is linear or *X* is constant (35; 36; 37; 38). The ratio function *f* (*π*_*N*_, *π*_*S*_) = *π*_*N*_ /*π*_*S*_ is convex in *π*_*S*_ for fixed *π*_*N*_ *>* 0, therefore, 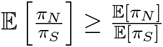. That is, the mean of ratios is greater than or equal to the ratio of means. For a set of *G* genes,

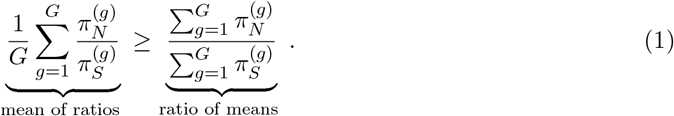

These two quantities use different weighting schemes. The mean of ratios, 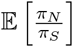, weights all genes equally, giving the same influence to a gene with few polymorphisms as one with many. These genes with few segregating sites are more susceptible to spurious results because of high sampling variance. The ratio of means, 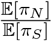, pools all variants genome-wide, therefore genes with more polymorphisms naturally contribute more, providing a precision-weighted estimate of genome-wide selection pressure.

The magnitude of the discrepancy between the two metrics depends strongly on the coefficient of variation (ℂ 𝕍) of *π*_*S*_: 𝕍 𝕍 (*π*_*S*_) = *σ*_*πS*_ /*𝔼* [*π*_*S*_]. When?𝔻??𝔻? is large (high variance in *π*_*S*_ relative to its mean), genes with low *π*_*S*_ dominate the mean of ratios because 1/*π*_*S*_ is large for these genes, and these genes often have high estimated *π*_*N*_ /*π*_*S*_ due to sampling variance. Indeed, empirically, the level of inflation between the mean of ratios and ratio of means correlates strongly with ℂ 𝕍 (*π*_*S*_) (Spearman *ρ* = 0.89, *p* < 0.001) and with the proportion of genes having zero synonymous diversity (*ρ* = 0.84, *p* < 0.001) across populations.

Next, I consider the contributions of sampling variance to patterns of gene-level *π*_*N*_ /*π*_*S*_. The variance of *π*_*N*_ /*π*_*S*_ can be approximated using the delta method (39), I have

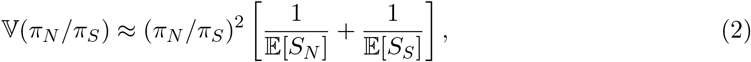

where *S*_*N*_ and *S*_*S*_ are the number of nonsynonymous and synonymous segregating sites, respectively. The corresponding coefficient of variation is

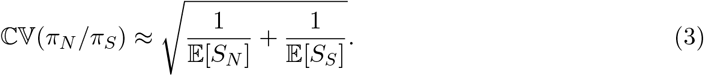

When both 𝔼 [*S*_*N*_] and 𝔼 [*S*_*S*_] are small (as occurs in populations with small *N*_*e*_), the coefficient of variation can exceed 1, meaning the standard deviation is larger than the mean. In this regime, single-gene *π*_*N*_ /*π*_*S*_ estimates are imprecise, largely governed by whether particular segregating sites happen to occur or not.

I applied the approximation from eqs. (2)-(3) to determine the expected proportion of genes exceeding *π*_*N*_ /*π*_*S*_ = 1 under neutral sampling variance. Using the genome-wide ratio of *π*_*N*_ /*π*_*S*_ = 0.56, assuming 𝔼 [*S*_*N*_] = 0.56 *×* 𝔼 [*S*_*S*_], the standard deviation *σ* = ℂ 𝕍 *×* 0.56 for genes with varying numbers of synonymous segregating sites. Under a normal approximation, I computed the probability that a gene’s *π*_*N*_ /*π*_*S*_ ratio exceeds 1 purely by sampling variance. This null expectation provides a baseline for identifying genes with genuine signals of relaxed constraint or positive selection beyond stochastic variation.

Table 1 shows that 25−32% of genes are expected to have *π*_*N*_ /*π*_*S*_ *>* 1 under neutrality when the number of synonymous segregating sites is very low. Comparatively, only 3 − 7% of genes with 10- 15 *S*_*S*_ should exceed *π*_*N*_ /*π*_*S*_ = 1 by chance under neutral sampling variance. These are calculated assuming genome-wide *π*_*N*_ /*π*_*S*_ = 0.56 and eqs. (2)-(3). Genes that consistently exceed this threshold despite having sufficient diversity likely experience genuine evolutionary forces beyond purifying selection. In the genomic data, when stratified by segregating sites, genes with <10 segregating sites show mean *π*_*N*_ /*π*_*S*_ of 2.9–17.2 across populations, whereas genes with *>*50 sites yield stable estimates (0.55–0.75) approaching genome-wide values. This segregating site dependence matches predictions from sampling variance theory, produced by the excess of rare variants predicted by a Beta coalescent. Consequently, 30–52% of genes show *π*_*N*_ /*π*_*S*_ *>* 1 across populations (mean = 40%), with 70–90% of low-segregating-site count genes but only 10–20% of high-segregating-site count genes exceeding this threshold.

**Table 1:** Expected proportion of genes exceeding *π*_*N*_ /*π*_*S*_ = 1 by sampling variance alone under the lifecycle model. Calculations assume genome-wide *π*_*N*_ /*π*_*S*_ = 0.56, and calculate *S*_*N*_ = 0.56*S*_*S*_ as an approximation. CV and *σ* calculated using eqs. 2 & 3. The z-score represents how many standard deviations above the genome-wide mean a value of *π*_*N*_ /*π*_*S*_ = 1 sits for genes with a given number of segregating sites assuming a normal distribution.

| $S_S$ | $S_N$ | CV | $\sigma$ | z-score | $P(\pi_N/\pi_S > 1)$ |
| --- | --- | --- | --- | --- | --- |
| 1 | 0.6 | 1.67 | 0.94 | 0.47 | 31.9% |
| 2 | 1.1 | 1.18 | 0.66 | 0.67 | 25.1% |
| 5 | 2.8 | 0.75 | 0.42 | 1.05 | 14.7% |
| 10 | 5.6 | 0.53 | 0.30 | 1.49 | 6.8% |
| 15 | 8.4 | 0.43 | 0.24 | 1.82 | 3.4% |
| 20 | 11.2 | 0.37 | 0.21 | 2.11 | 1.7% |
| 25 | 14.0 | 0.33 | 0.19 | 2.37 | 1.0% |
| 30 | 16.8 | 0.31 | 0.17 | 2.59 | 0.5% |

To empirically assess the uncertainty in gene-level *π*_*N*_ /*π*_*S*_ estimates, I performed bootstrapping to get confidence intervals for 4,299 genes with *S >* 0 in the Ghana population sample (Fig. 4). For each gene, I resampled segregating sites 1,000 times using Poisson resampling (independently for *S*_*N*_ and *S*_*S*_), calculating the 95% confidence interval from the bootstrap distribution. Fig. 4A shows that confidence intervals scale with the number of segregating sites as CI width ∝ *S*^−0.61^, closely matching the theoretical prediction of *S*^−0.5^. The observed coefficient of variation tracks remarkably well with the theoretical prediction from eq. (3) (Fig. 4B). Ghana is given here as a representative example, with other countries showing qualitatively similar patterns.

**Figure 4.**
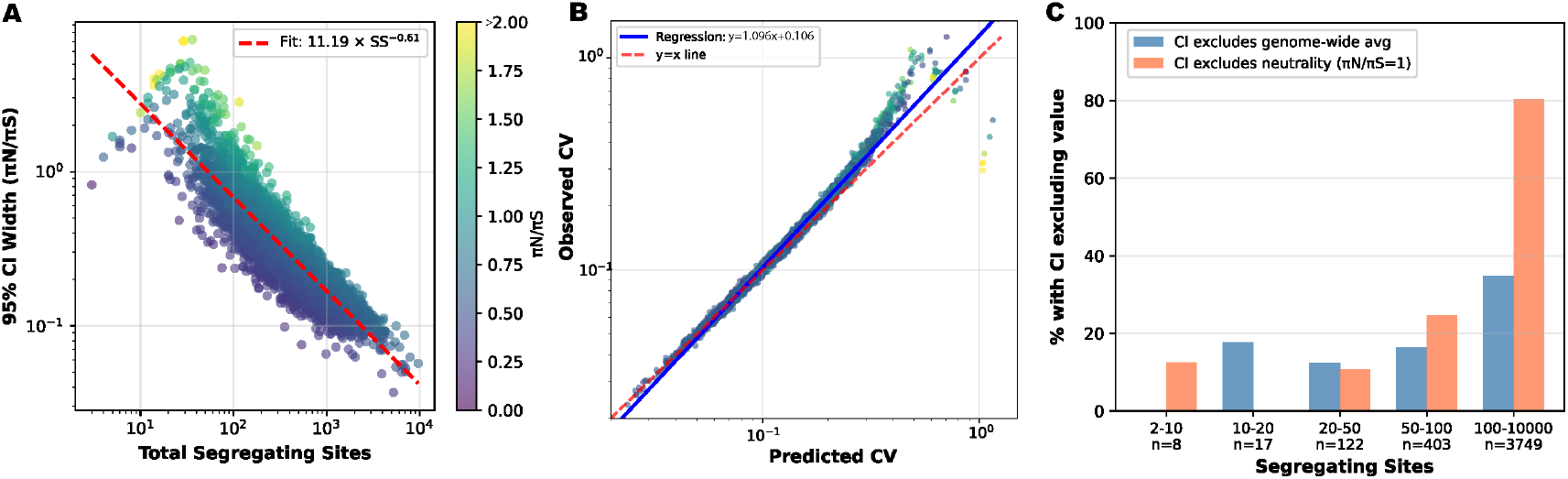
Bootstrapping shows high uncertainty gene-level *π*_*N*_ /*π*_*S*_ driven by low numbers of segregating sites. Bootstrap 95% confidence intervals (CIs) for gene-level *π*_*N*_ /*π*_*S*_ ratios in Ghana (*n*=4,299 genes with ≥1 segregating site). **(A)** CI width decreases with total segregating sites following an approximate power law, colored by observed *π*_*N*_ /*π*_*S*_ ratio. **(B)** Observed coefficient of variation (ℂ 𝕍) from bootstrap resampling scales with the Poisson-based theoretical prediction from eq. (3) in which ℂ 𝕍 decreases with increasing number of segregating sites. **(C)** The percent of genes for which their 95% confidence interval does not include genome-wide average (*π*_*N*_ /*π*_*S*_ = 0.57, blue) or neutrality (*π*_*N*_ /*π*_*S*_ = 1, orange) binned by the number of segregating sites per gene. This is a proxy for statistical power to detect deviation, which increases with segregating site counts. Results were highly consistent across countries.

Whereas the theoretical formula predicts the magnitude of uncertainty, Fig. 4C considers statistical power to detect deviations from neutral expectations. Fig. 4C plots the percentage of genes whose 95% confidence intervals exclude either the genome-wide average (*π*_*N*_ /*π*_*S*_ = 0.57) or neutrality (*π*_*N*_ /*π*_*S*_ = 1). For genes in the lowest segregating site bins (*S* = 2–10 and *S* = 10–20), fewer than 20% have confidence intervals excluding the genome-wide average, despite many having point estimates that appear elevated or depressed. Similarly, only 12% and 10% of genes in these bins have CIs excluding neutrality. This demonstrates that most apparent signals in low-diversity genes are statistically indistinguishable from neutral expectations. As segregating site counts increase, statistical power improves dramatically.

In sum, apparent elevation of *π*_*N*_ /*π*_*S*_ in low-diversity genes reflects sampling noise amplified by properties of ratio statistics rather than signals of selection. Further properties of *π*_*N*_ /*π*_*S*_ are discussed in Supplementary Methods, including skew from zero-truncation.

### 4.2 Rare variants drive noisy estimates and mask selection

Rare variants, which dominate the number of segregating sites under a Beta coalescent, contribute disproportionately to sampling variance in ratio estimates, especially as their per-locus impact on pairwise diversity is minimal. To demonstrate the various impacts of biological mechanisms (rare variants), and statistical properties (sampling variance) in low-diversity genes, I applied two complementary filters to the genomic data (Fig. 5A). Importantly, the majority of genes lack sufficient common variation for robust estimation (Fig. 5B).

**Figure 5.**
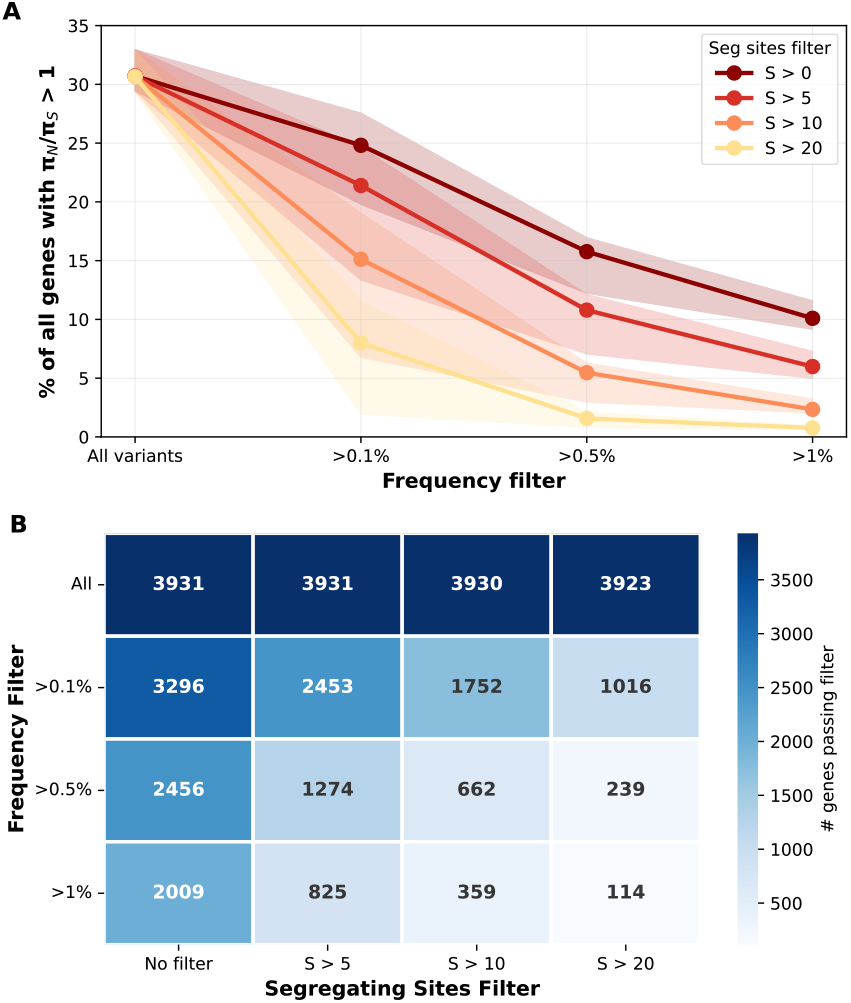
Filtering empirical data shows the combined impact of rare variants and number of segregating sites on the proportion of genes with *π*_*N*_ /*π*_*S*_ *>* 1. Lines show median values with shaded regions indicating interquartile range (25th–75th percentile) across countries. **(A)** Effect of frequency filtering for different segregating site thresholds. When rare variants are included ∼36% of genes show *π*_*N*_ /*π*_*S*_ *>* 1 regardless of minimum segregating site threshold. Frequency filtering substantially reduces apparent elevation across all segregating site thresholds. **(B)** Number of genes passing each filter combination. The proportion of genes within each bin with *π*_*N*_ /*π*_*S*_ *>* 1 stays roughly similar.

Filtering tests two related predictions, the first biological and the second statistical. First, filtering by frequency removes recent, rare among-host variants and therefore focuses on alleles that have the opportunity to be under selection among multiple hosts over time. Second, requiring an adequate number of segregating sites per gene should reduce sampling variance in ratio estimates.

The z-scores in Table 1 highlight why filtering by segregating site count may help to identify biological signals. The z-score represents how many standard deviations above the genome-wide mean (*π*_*N*_ /*π*_*S*_ = 0.56) a value of *π*_*N*_ /*π*_*S*_ = 1 sits. Genes with low numbers of segregating sites represent false positives driven by Jensen’s inequality and sampling noise rather than biological signal. Notably, Figs. 5 and S3 shows that the patterns predicted by the model shown in Table 1 only hold in the empirical data after filtering out rare variants, which perhaps haven’t been subject to selection pressures across the lifecycle or transmission drift yet. When all variants are included, the proportion of genes with *π*_*N*_ /*π*_*S*_ *>* 1 remains high and constant regardless of segregating site threshold, contrary to Table 1’s prediction that this proportion should decline sharply with increasing segregating sites. Only after frequency filtering does the expected segregating-site dependence emerge, suggesting that rare variants generated by Beta coalescence must first be removed before the sampling variance framework fits. These two processes—the excess rare variation from Beta coalescence, and overall low number of segregating sites—work together, but are perhaps expected to be of different level of importance in different geographic regions given different population-level diversity and *α*.

Next, I considered biological interpretation of the genes with *π*_*N*_ /*π*_*S*_ *>* 1 with and without rare variants using functional enrichment tests and comparisons across countries. The set of filtered genes without rare variants shows more interpretable signals of selection. Despite containing on average 3.2-fold fewer genes, for both filtered and unfiltered sets, the most robustly enriched processes across populations were consistently related to host-pathogen interaction. “Entry into host” (GO:0044409) was significantly enriched in 16/18 countries in the filtered analysis versus 14/18 in the unfiltered analysis. “Malaria pathogenesis” (GO:0020009) was enriched in 15 filtered and 18 unfiltered country datasets, while “host cell surface binding” (GO:0046812) showed enrichment in 15 filtered versus only 8 unfiltered analyses. That is, sets of genes that we expect to be under selection are retained through filtering, with the large proportion of genes lost contributing minimal true signal.

Analysis of geographic sharing identified 17 genes with *π*_*N*_ /*π*_*S*_ *>* 1 across all 18 countries in the filtered dataset (Table S6), a significant enrichment under a binomial test *p* < 10^−6^. These universally selected genes included well-characterized targets of immune pressure: apical membrane antigen 1 (*AMA1*), merozoite surface protein 6 (*MSP6*), Duffy binding-like merozoite surface protein (*DBLMSP*), and multiple members of the 6-cysteine protein and SURFIN families. In contrast, 84 genes were shared as *π*_*N*_ /*π*_*S*_ *>* 1 across the 18 countries in the unfiltered set, consistent with the substantially higher overall number of genes with *π*_*N*_ /*π*_*S*_ *>* 1. The list was partially overlapping but contained many unannotated or otherwise difficult to interpret genes as well, suggesting filtering improves the signal-to-noise ratio.

Finally, I considered the impact of filtering by testing for genes that changed whether they were above or below the *π*_*N*_ /*π*_*S*_ = 1 line. Across 18 countries, an average of 1,399 genes (range: 1,047–1,983) had *π*_*N*_ /*π*_*S*_ *>* 1 in the unfiltered dataset (Figure 5A). Applying frequency and site count filters (*>*0.5% frequency and *>*5 segregating sites per gene) reduced this to an average of 431 genes per country (range: 228–802), representing a 69% reduction in candidate genes. However, the filtered set was not simply a subset of the unfiltered genes. While 82.4% of filtered genes were present in the unfiltered set, 17.6% (mean: 86 genes per country, range: 5–151) gained *π*_*N*_ /*π*_*S*_ *>* 1 status only after filtering (Figure 5B).

These “gained” genes (*n* = 724 unique genes across all countries) often had elevated *π*_*N*_ /*π*_*S*_ ratios in the unfiltered analysis (mean: 0.85), which then increased after filtering (mean: 1.46). While most gained genes showed moderate increases in *π*_*N*_ /*π*_*S*_ (mean increase: 0.62), some showed dramatic shifts in a subset of countries, such as perforin-like protein 2 (*π*_*N*_ /*π*_*S*_: 0.76 → 6.36) and BRO1 domain-containing protein (*π*_*N*_ /*π*_*S*_: 0.87 → 11.40), where filtering revealed strong selection signals that were masked by rare variants in the unfiltered analysis. Notably, gained genes included well-characterized malaria antigens such as merozoite surface protein 1 (MSP1, *π*_*N*_ /*π*_*S*_: 0.97 → 1.01), merozoite organizing protein (MOP, *π*_*N*_ /*π*_*S*_: 0.72 → 1.21), sporozoite surface antigen MB2 (*π*_*N*_ /*π*_*S*_: 0.93 → 2.37), rhoptry-associated protein (RAP, *π*_*N*_ /*π*_*S*_: 0.76 → 1.96), Female development protein FD1 (*π*_*N*_ /*π*_*S*_: 0.76 → 1.98), and WD repeat-containing protein (*π*_*N*_ /*π*_*S*_: 0.84 → 2.44), demonstrating that variant filtering reveals genuine selection targets obscured by noisy rare variants in unfiltered analysis.

Enrichment analysis of the 724 genes pooled across all populations revealed by filtering but missed in unfiltered analysis showed significant enrichment for host-pathogen interaction functions. These genes included surface and exported proteins, invasion-related proteins, and sexual stage proteins. GO term analysis showed enrichment for “entry into host,” “Maurer’s cleft,” “cell surface,” and “malaria pathogenesis”. The recovery of genes putatively under selection through variant filtering demonstrates that this approach is essential for comprehensive identification of selection targets, not merely a method to reduce noise.

In sum, the filtering analysis presented here is not intended as a universal recommendation for how to analyze malaria genomic data, but rather as a diagnostic tool that guides interpretation of *π*_*N*_ /*π*_*S*_ ratios in this system. In systems well-represented by the standard Kingman coalescent the set of genes with *π*_*N*_ /*π*_*S*_ *>* 1 is enriched for genes under positive or balancing selection because the genealogy generates relatively few rare variants. The ∼ 20 − 40% of *P. falciparum* genes exceeding *π*_*N*_ /*π*_*S*_ *>* 1 might therefore seem to indicate widespread adaptive evolution or relaxed constraint. Rather, the filtering analysis supports the proposal suggesting most of genes with *π*_*N*_ /*π*_*S*_ *>* 1 are elevated because of a combination of (1) a substantial proportion of segregating sites being low-information contributors to *π* because they are recent and rare, and therefore have not yet been tested by selection across multiple hosts and transmission cycles, and (2) high sampling variance from a small number of segregating sites per gene, which is most relevant in Southeast Asia.

## 5 Conclusions

Here, I suggest that *Plasmodium falciparum* genealogies are better represented as following a multiple-merger Beta coalescent reflecting extreme reproductive skew, and that this likely genealogical structure systematically biases standard population genetic statistics, including estimates of Tajima’s *D* and estimators of *N*_*e*_. Further, the rare variant excess generated by multiple-merge coalescence interacts with the mathematical properties of ratio statistics to inflate gene-level *π*_*N*_ /*π*_*S*_ estimates, while filtering reveals hundreds of previously masked selection targets including well-characterized antigens.

The two sources of inflation—rare variant excess from Beta coalescence and sampling variance from low segregating sites—differ in their relative importance between geographic regions. African populations exhibit stronger multiple-merger signatures (*α* ≈ 1.36) but higher absolute diversity (mean *π*_*S*_ ≈ 0.0006), whereas Southeast Asian populations show weaker Beta coalescent signatures (*α* ≈ 1.82) but lower diversity (mean *π*_*S*_ ≈ 0.0003–0.0004). In Africa, the rare variant excess is the primary driver of inflation (mean of ratios 1.5–2.0-fold higher than ratio of means). In Southeast Asia, low diversity amplifies sampling variance, producing more extreme inflation (5.9–8.6-fold higher) despite weaker multiple-merger dynamics. This geographic difference manifests in the proportion of genes excluded due to *π*_*S*_ = 0 (16–30% in Southeast Asia versus 1–4% in Africa) and in the proportion showing *π*_*N*_ /*π*_*S*_ *>* 1 before filtering. After filtering addressing both the number of segregating sites and variant frequency, geographic differences in apparent elevation are substantially reduced.

Other processes may cause an excess of rare variants, including population expansion or background selection. I explicitly compared the fit of expansion models, finding a better fit for the U-shaped Beta coalescent models. Henry et al. (2025) (10) estimated the expected reduction in diversity from BGS in *P. falciparum* to be minimal, owing to short chromosomes and high recombination rates.

This fit over a traditional Kingman coalescent does not provide a specific mechanism, but rather emphasizes the empirically-observed shape of the SFS, particularly the excess very-rare variants. Several features of malaria transmission biology could generate such heavy-tailed offspring distributions that produce genealogies consistent with a Beta coalescent. First, for many pathogens transmission success is overdispersed, perhaps owing to heterogeneities in biting rate, network structure, and gametocyte density (40; 41; 42). Second, the lifecycle concentrates ancestry into a small number of highly successful transmission events per generation, the offspring distribution at the between-host level is expected to be heavily right-skewed. Particularly if multiplicity of infection is high, when only ∼10–100 parasites survive the bottleneck, stochastic loss or amplification of lineages is likely. High multiplicity of infection also increases co-transmission of related lineages into mosquitoes where they may recombine. Finally, population-level changes like rapid adaptation or seasonal population size fluctuations may contribute.

The statistical challenges of common population-genetic summary statistics described here are not unique to malaria. Similar biases may arise in any system where demographic or life-history processes generate multiple-merger coalescent dynamics: pathogens with transmission bottlenecks (*Mycobacterium tuberculosis*, influenza, SARS-CoV-2 superspreader events), organisms with extreme fecundity (marine invertebrates, many fungi), and any system where small *N*_*e*_ produces abundant rare variants (22; 23; 43; 21; 44; 45). Durrett (2013) (46) considered analogous difficulties distinguishing driver from passenger mutations in cancer genomics, where nonsynonymous to synonymous ratios lack power when genes have few mutations. The mathematical principle connecting these systems is general: ratio statistics with low-diversity denominators are dominated by sampling variance rather than biological signal.

Analogous issues have been noted for other ratio statistics. For *F*_*ST*_ estimation, computing the statistic as an average of per-locus ratios produces biased estimates compared to computing it as a ratio of pooled numerators and denominators (47; 48). In comparisons of closely related species, low *d*_*S*_ values rather than elevated *d*_*N*_ often drive apparently elevated *d*_*N*_ /*d*_*S*_ ratios (49). The principle is consistent: weighting all loci equally gives disproportionate influence to low-diversity loci where sampling variance dominates, while pooling provides precision-weighted estimates.

Several limitations constrain interpretation. The Beta coalescent fit is empirical rather than derived from explicit epidemiological transmission models; mechanistic integration of transmission dynamics, recombination, and multiplicity of infection remains an important future direction. Filtering thresholds applied here are heuristic, and optimal approaches may vary across populations with different transmission intensities.

Looking forward, this framework suggests that genealogical models explicitly accommodating multiple-merger dynamics should be considered rather than Kingman-based approaches for pathogen genomic inference. More broadly, before interpreting genomic signatures of selection in any high-fecundity or high-skew system, we must first understand the genealogical structure that generated the data.

## Materials and Methods

### Genomic data and filtering

Whole-genome sequence data were obtained from the MalariaGEN *Pf* 8 release, which provides variant calls for *Plasmodium falciparum* samples collected globally. We analyzed 18 populations with at least 200 samples passing quality control: 10 African countries (Cameroon, Democratic Republic of Congo (DRC), Gambia, Ghana, Kenya, Malawi, Mali, Mozambique, Nigeria, Tanzania) and 8 Asian/Oceanian countries (Bangladesh, Cambodia, India, Laos, Myanmar, Papua New Guinea (PNG), Thailand, Vietnam). Sample metadata and quality control flags followed designation from the `malariagen_data` Python package (v8.0). Only samples flagged as “QC pass = True” in the MalariaGEN pipeline were retained, which filters for adequate sequencing coverage, low contamination rates, and successful variant calling. Variants were restricted to those passing MalariaGEN’s variant-level quality filters (`variant_filter_pass = True`), which removes sites with evidence of systematic genotyping errors, excessive missingness, or strand bias. Analyses were restricted to the 14 nuclear chromosomes.

### Gene annotation and variant classification

Gene annotations were obtained from PlasmoDB release 68. Coding sequences were extracted for all protein-coding genes. Specifically, for each position within each codon, I counted the number of possible substitutions (to A, T, C, or G) resulting in the same amino acid (synonymous) versus a different amino acid (nonsynonymous), weighting each site by the proportion of changes in each category. Due to extreme codon usage bias in the AT-rich *P. falciparum* genome (∼80% AT content), this yields over 4 nonsynonymous sites per synonymous site, substantially higher than the ∼2.5–3 ratio typical of organisms with balanced nucleotide composition.

Callable sites per gene were calculated as total sites multiplied by the variant pass rate (proportion of variants within the gene passing quality filters). Genes with pass rates below 70% were excluded from downstream analyses, yielding 4,310 genes for analysis.

For each SNP, the reference codon was extracted from the *P. falciparum* 3D7 reference genome (accessed via `malariagen_data`). The alternate codon was constructed by substituting the alternate allele at the appropriate codon position, and amino acid translations were compared using the standard genetic code. Variants were classified as synonymous if reference and alternate amino acids were identical, nonsynonymous if they differed, and noncoding if outside annotated CDS regions.

### Nucleotide diversity

Nucleotide diversity (*π*) was calculated as the average pairwise differences per site. For each segregating site, heterozygosity was computed as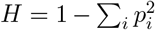, where *p*_*i*_ is the frequency of allele *i* among non-missing calls. A finite-sample correction was applied: 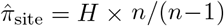 where *n* is the number of successfully called alleles at that site. Gene-level *π* was calculated by summing corrected per-site heterozygosity values and dividing by callable sites. Genome-wide diversity was computed as the site-weighted sum across genes: *π*_genome_ = ∑_*g*_ (*π*_*g*_ *× L*_*g*_)/ ∑_*g*_ *L*_*g*_, where *L*_*g*_ represents callable sites for gene *g*. Separate calculations were performed for synonymous (*π*_*S*_) and nonsynonymous (*π*_*N*_) sites.

Bootstrap 95% confidence intervals were generated using 1,000 Poisson resampling replicates of segregating sites per gene (independently for *S*_*N*_ and *S*_*S*_).

### Empirical inference of *N*_*e*_

Diversity-based effective population size was estimated as 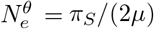 for haploid blood-stage parasites, using mutation rate *µ* = 1.5 *×* 10^−9^ per site per generation. This approach implicitly weights coalescence times toward phases where lineages can coalesce, particularly the within-host expansion phase.

LD-based effective population size was estimated by fitting the decay of *r*^2^ with physical distance to the expectation for haploid organisms with inbreeding,

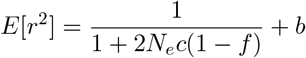

where *c* is recombination rate (7.4 *×* 10^−7^ per bp per generation), *f* is the inbreeding coefficient representing the probability of identity by descent due to selfing, and *b* captures background LD from factors other than drift. Region-specific inbreeding coefficients were applied as heuristic proxies for transmission intensity: *f* = 0.3 for sub-Saharan African populations (higher transmission, more outcrossing during mosquito stage) and *f* = 0.7 for Asian and Oceanian populations (lower transmission, higher selfing rates). These inbreeding rates are representative rather than exact estimates of *f*, and are used to represent general trends; considering the whole range of *f* ∈ [0, 1], the qualitative patterns of *N*_*e*_-LD <*N*_*e*_-theta hold. Parameters were estimated by nonlinear least squares fitting.

### Site frequency spectrum and Beta coalescent model fitting

SFS were calculated for each population using all biallelic SNPs on the 14 nuclear chromosomes passing MalariaGEN quality filters. To ensure comparable sample sizes across populations, each population was subsampled to *n* = 200 individuals before SFS calculation. For each SNP, derived (alternate) allele counts were computed from haploid genotypes, excluding missing calls. The unfolded SFS was constructed by tabulating the number of SNPs at each derived allele count. Fourfold degenerate positions were identified by iterating through all CDS features and extracting third codon positions where any nucleotide substitution is synonymous.

Beta(2 − *α, α*) coalescent models were fit independent for each population to the observed SFS by maximum likelihood. Under this model, the expected proportion of SNPs at frequency *k* follows

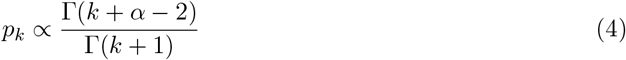

where Γ denotes the gamma function. The parameter *α* was estimated by maximizing the loglikelihood, given by

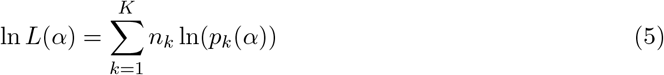

where *n*_*k*_ is the observed count of SNPs at frequency *k*. Optimization used bounded scalar minimization over *α* ∈ (1.05, 1.95), fitting to the first *K* = 100 frequency classes. Model fit was assessed using Pearson correlation between log-transformed observed and expected frequency spectra.

Model comparison used likelihood ratio tests (LRT),

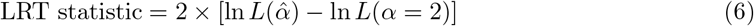

comparing the fitted Beta coalescent to the null Kingman coalescent (*α* = 2.0). Under the null hypothesis, the LRT statistic follows a *χ*^2^ distribution with 1 degree of freedom. All 18 populations rejected the Kingman coalescent with *p* < 0.001.

For main text analyses, the SFS from all variants was used. Replication with only fourfold degenerate sites is in the Supplement and is highly consistent (Pearson correlation, r = 0.997, and a mean *α* difference by country between the full-variant and fourfold-degenerate fits of only −0.005).

**Tajima’s** *D* Tajima’s *D* was calculated using the scikit-allel implementation (v1.3.5), computing the standardized difference between mean pairwise diversity (*θ*_*π*_) and Watterson’s estimator (*θ*_*W*_).

### Gene ontology enrichment

Functional enrichment used Gene Ontology (GO) annotations from PlasmoDB release 68. The background gene set consisted of all 4,310 protein-coding genes with *π*_*S*_≠ 0 in the filtered dataset. For each population, the set of genes with *π*_*N*_ /*π*_*S*_ *>* 1 after filtering (*>*0.5% frequency, *>*5 segregating sites) were tested. Enrichment was assessed using hypergeometric tests and p-values follow Benjamini-Hochberg FDR *q* < 0.05.

## Acknowledgments

This paper arose out of a meeting hosted by the Society for Modeling and Theory in Population Biology at the National Institute for Theory and Mathematics in Biology, particularly from conversations with John Wakeley, Katia Koelle, and Maria Orive. I also thank Nandita Garud and Joshua Schraiber for useful discussion. Funding by National Institutes of Health R35-GM133481 and R01-AI175622 to AG.

## Data Availability

MalariaGEN *Pf* 8 data are publicly available through the MalariaGEN resource center, https://www.malariagen.net/data.

Analysis scripts are available at https://github.com/agoldberglab/Malaria_Beta_Coalescent.

