## Supplemental Materials for "Rare variation in malaria parasites biases population-genetic inference"

Amy Goldberg

Department of Human Genetics, University of California, Los Angeles, USA

#### 1 Fitting SFS models

I compared two coalescent models for the observed site frequency spectra: the Beta$(2-\alpha, \alpha)$ coalescent with multiple mergers and exponential population growth. Under the Beta coalescent, the expected SFS follows the asymptotic form

$$\mathbb{E}[\xi_k] \propto \frac{\Gamma(k+\alpha-2)}{\Gamma(k+1)} \quad (1)$$

which for large $k$ behaves as $k^{\alpha-3}$, producing a linear relationship on log-log axes with slope $(\alpha-3)$ for $\alpha \in (1,2)$ (1; 2). This approximation captures the power-law decay characteristic of multiple-merger genealogies and is most accurate at low-to-intermediate frequencies where the vast majority of segregating sites occur. For exponential growth with scaled rate $\rho$, the expected SFS takes the form

$$\mathbb{E}[\xi_k] \propto \frac{1}{k} \cdot \exp\left(-\frac{k}{\kappa}\right) \quad (2)$$

where $\kappa$ depends on sample size $n$ and growth rate following Polanski et al. (2003) (3). A key theoretical distinction between these models is their predicted shape on a log-log scale. Both Kingman and Beta coalescent produce linear SFS with slopes of $-1$ and $(\alpha-3)$ respectively, whereas exponential growth produces a curved SFS.

I estimated model parameters by maximum likelihood using all $K = n-1 = 199$ frequency classes of each population's unfolded SFS. Following Eldon et al. (2015) (2), I used the Poisson Random Field approximation, which treats frequency classes as independent Poisson random variables (4). This yields an approximate log-likelihood

$$\ell(\Pi) = \sum_{k=1}^{K} \xi_k \log \phi_k(\Pi) \quad (3)$$

where $\xi_k$ is the observed count at frequency $k$ and $\phi_k(\Pi) = \mathbb{E}[\xi_k]/\sum_j \mathbb{E}[\xi_j]$ is the normalized expected proportion under model $\Pi$. I maximized this likelihood over $\alpha \in (1.01, 1.99)$ for the Beta coalescent and $\rho \in (0.1, 10^5)$ for exponential growth using Brent's method.

For model comparison, I computed the log likelihood ratio $\Lambda_{\beta/G} = \ell(\hat{\alpha}) - \ell(\hat{\rho})$, where positive values favor the Beta coalescent. I also computed the minimum $\ell_2$ distance between the observed normalized SFS ($\zeta_k = \xi_k/S$) and the expected SFS under each model, following Equation 15 of Eldon et al. (2015) (2),

$$d_\beta = \inf_\alpha \sqrt{\sum_k (\phi_k(\alpha) - \zeta_k)^2}, \quad d_G = \inf_\rho \sqrt{\sum_k (\phi_k(\rho) - \zeta_k)^2} \quad (4)$$

A ratio $d_G/d_\beta > 1$ indicates the Beta coalescent provides a closer fit to the observed data.

The Beta coalescent provided a better fit than exponential growth by the likelihood ratio criterion in all 18 populations (Table S2). Log likelihood ratios ranged from +878 (Laos) to +51,389 (Mali). The fitted Beta coalescent parameter $\hat{\alpha}$ showed a regional gradient consistent with transmission intensity: African populations had the lowest values (mean $\hat{\alpha} = 1.36 \pm 0.04$), followed by South Asia ($1.56 \pm 0.00$), Southeast Asia ($1.82 \pm 0.03$), and Oceania (1.84).

For the two countries with "mixed" evidence for Beta coalescent over Kingman with growth, it stems from differences in the calculation of comparison statistics. The log-likelihood weighs each SNP equally, so the fit is driven by the most abundant frequency class (singletons, comprising 21–72% of SNPs). The $\ell_2$ distance weights each frequency class equally regardless of SNP count, giving more influence to intermediate and high-frequency variants where Beta and growth models can diverge. To make SFS more comparable across populations, all were built with each population down-sampled to $n = 200$ individuals. As a test, I fit $\alpha$ for $n = 1,000$ for Ghana and Vietnam, and results were highly similar, with a difference of 0.02 and 0.03, respectively.

### 2 Right-skew artifacts in ratio estimates.

Though likely more minor, multiple other mathematical artifacts may also create right-skewed ratio distributions for $\pi_N/\pi_S$.

First, when the relationship between $\pi_N$ and $\pi_S$ has a positive intercept, that is, genes with essentially zero synonymous diversity maintain some baseline nonsynonymous diversity, then ratios become mechanically inflated as $\pi_S$ approaches zero. If $\pi_N = a + b\pi_S + \epsilon$ where $a > 0$, then $\pi_N/\pi_S = a/\pi_S + b + \epsilon/\pi_S$, and the term $a/\pi_S$ creates an inverse relationship between $\pi_S$ and $\pi_N/\pi_S$ (5; 6). This introduces heteroscedasticity where $\mathrm{Var}(\pi_N/\pi_S) \approx a^2/\pi_S^2$, so uncertainty increases dramatically as $\pi_S$ decreases. Therefore, the right tail is amplified; though most genes cluster near the genome-wide average, genes with low $\pi_S$ exhibit much wider variation, with rare extreme values driving the long right tail of the distribution.

I compared linear regression of $\pi_N$ on $\pi_S$ for each country for the set of all variants to the same regression for the filtered set of variants occurring at $\geq 0.5\%$ frequency (per population). Regression of $\pi_N$ on $\pi_S$ across 18 global *P. falciparum* populations reveals universal positive intercepts when using all variants (mean $a = 1.4 \times 10^{-4}$), indicating that genes with very low synonymous diversity maintain a baseline level of nonsynonymous diversity. However, filtering to common variants ($>$ 0.5% frequency) reduces these intercepts by 62% (to mean $a = 5.25 \times 10^{-5}$), with three low-diversity populations (Vietnam, Cambodia, Thailand) exhibiting negative intercepts after filtering, and improves regression model fit ($R^2$ increases from 0.18 to 0.29). This suggests that positive intercepts are primarily driven by rare variants in low-diversity genes, where sampling variance and weakly deleterious mutations create apparent "floors" in $\pi_N$.

Second, zero-truncation amplifies right skew. Both $\pi_N$ and $\pi_S$ are always greater than or equal to zero. Therefore, sampling variance that would naturally produce some negative estimates instead creates a pile-up of small positive values near zero. For genes with very low true diversity, this produces a right-skewed distribution of ratio estimates.

Indeed Chang et al. (2013) (7) noted that $\pi_S$ was reduced rather than $\pi_N$ increased for the subset of genes with $\pi_N/\pi_S > 1$, suggesting that these right-skew effects greatly impact the proportion of genes with $\pi_N/\pi_S > 1$ that are of interest. This pattern partially holds in the data analyzed here, where comparing the set of genes with $\pi_N/\pi_S > 1$ to genes with $\pi_N/\pi_S \leq 1$ in each of the 18 populations, on average $\pi_S$ is 4.5-times smaller ($8.43 \times 10^{-5}$ compared to $3.83 \times 10^{-4}$) whereas $\pi_N$ is 2.2-times larger ($2.60 \times 10^{-4}$ compared to $1.17 \times 10^{-4}$) in the set of genes with $\pi_N/\pi_S > 1$.

#### Supplementary Tables and Figures

| **Population** | **Sample Size** | **Total SNPs** | **Region** |
|---|---|---|---|
| Cameroon | 294 | 95,177 | Africa |
| DRC | 1,549 | 105,530 | Africa |
| Gambia | 1,998 | 102,899 | Africa |
| Ghana | 2,000 | 102,708 | Africa |
| Kenya | 2,142 | 90,583 | Africa |
| Malawi | 681 | 86,089 | Africa |
| Mali | 2,428 | 111,455 | Africa |
| Mozambique | 1,348 | 95,135 | Africa |
| Nigeria | 1,303 | 104,516 | Africa |
| Tanzania | 1,144 | 97,772 | Africa |
| PNG | 251 | 38,669 | Oceania |
| Bangladesh | 1,658 | 74,009 | South Asia |
| India | 318 | 65,200 | South Asia |
| Cambodia | 2,282 | 33,265 | Southeast Asia |
| Laos | 1,994 | 35,842 | Southeast Asia |
| Myanmar | 1,268 | 38,283 | Southeast Asia |
| Thailand | 1,157 | 32,942 | Southeast Asia |
| Vietnam | 2,700 | 28,960 | Southeast Asia |

Table 1: Summary of 18 *Plasmodium falciparum* populations from MalariaGEN *Pf8*. For each population, we report the sample size (number of parasites sequenced), total number of SNPs, and geographic region. The full dataset for Ghana has 6,653 individuals, but it was downsampled to 2,000 to be in line with other populations.

| Population | $\alpha$ | Singletons | $\Delta\text{AIC}_{\text{Kingman}}$ | $\Delta\text{AIC}_{\text{Growth}}$ | $d_G/d_\beta$ | Best Model |
|---|---|---|---|---|---|---|
| Ghana | 1.30 | 72.4% | 165,111 | 95,228 | 24.6 | Beta |
| Mali | 1.31 | 72.2% | 178,794 | 102,778 | 26.3 | Beta |
| DRC | 1.34 | 68.1% | 155,372 | 87,054 | 25.1 | Beta |
| Nigeria | 1.35 | 67.3% | 151,076 | 80,509 | 19.2 | Beta |
| Tanzania | 1.37 | 64.7% | 131,854 | 73,368 | 18.0 | Beta |
| Malawi | 1.38 | 64.2% | 113,785 | 65,240 | 17.0 | Beta |
| Cameroon | 1.39 | 62.8% | 123,615 | 66,363 | 10.0 | Beta |
| Mozambique | 1.39 | 62.2% | 123,272 | 64,719 | 10.5 | Beta |
| Kenya | 1.41 | 60.6% | 110,961 | 58,347 | 10.2 | Beta |
| Gambia | 1.41 | 61.3% | 124,647 | 64,965 | 28.4 | Beta |
| Bangladesh | 1.56 | 44.2% | 52,910 | 27,728 | 1.1 | Beta |
| India | 1.56 | 44.2% | 46,342 | 22,851 | 1.3 | Beta |
| Vietnam | 1.82 | 30.4% | 3,217 | 4,678 | 2.0 | Beta |
| Cambodia | 1.78 | 33.2% | 5,856 | 5,454 | 3.2 | Beta |
| Myanmar | 1.81 | 29.0% | 5,150 | 3,608 | 3.2 | Beta |
| Thailand | 1.83 | 27.9% | 3,517 | 2,671 | 3.3 | Beta |
| Laos | 1.86 | 21.4% | 2,660 | 1,855 | 0.8 | Mixed$^\dagger$ |
| PNG | 1.84 | 23.0% | 3,608 | 2,087 | 0.6 | Mixed$^\dagger$ |

Table 2: Beta coalescent parameter estimates and model comparison for 18 *Plasmodium falciparum* populations. $\alpha$ is the Beta coalescent parameter (smaller values indicate stronger multiple-merger signal); populations are sorted by $\alpha$ from lowest to highest. Singleton proportion is also shown. $\Delta\text{AIC} = 2(\ell_{\text{Beta}} - \ell_{\text{alt}}) - 2(k_{\text{Beta}} - k_{\text{alt}})$, where $\ell$ is log-likelihood and $k$ is number of parameters; since Beta and Growth each have one parameter, $\Delta\text{AIC}_{\text{Growth}} = 2(\ell_{\text{Beta}} - \ell_{\text{Growth}})$. $d_G/d_\beta$ is the ratio of minimum $\ell_2$ distances between observed and expected SFS (Eldon et al. 2015), with values $> 1$ favoring Beta. The log-likelihood weights each SNP equally, so fit is dominated by singletons (21–72% of SNPs); the $\ell_2$ distance weights each frequency class equally, giving more influence to intermediate and high-frequency variants.

$^\dagger$Likelihood favors Beta, but $\ell_2$ distance favors Growth.

| Country | $D_S$ (mean ± SD) | $D_N$ (mean ± SD) | Genome-wide $D$ |
|---|---|---|---|
| Bangladesh | $-1.41 \pm 0.51$ | $-1.69 \pm 0.52$ | $-2.14$ |
| Cambodia | $-1.41 \pm 0.47$ | $-1.86 \pm 0.48$ | $-1.21$ |
| Cameroon | $-1.68 \pm 0.50$ | $-1.86 \pm 0.55$ | $-2.45$ |
| DRC | $-1.95 \pm 0.43$ | $-2.21 \pm 0.41$ | $-2.51$ |
| Gambia | $-1.90 \pm 0.45$ | $-2.19 \pm 0.40$ | $-2.42$ |
| Ghana | $-2.12 \pm 0.40$ | $-2.37 \pm 0.33$ | $-2.55$ |
| India | $-1.56 \pm 0.50$ | $-1.90 \pm 0.51$ | $-2.12$ |
| Kenya | $-1.79 \pm 0.46$ | $-2.02 \pm 0.46$ | $-2.42$ |
| Laos | $-1.30 \pm 0.52$ | $-1.72 \pm 0.51$ | $-1.16$ |
| Malawi | $-1.74 \pm 0.47$ | $-1.94 \pm 0.51$ | $-2.46$ |
| Mali | $-2.04 \pm 0.43$ | $-2.31 \pm 0.37$ | $-2.55$ |
| Mozambique | $-1.87 \pm 0.43$ | $-2.19 \pm 0.40$ | $-2.46$ |
| Myanmar | $-1.27 \pm 0.58$ | $-1.59 \pm 0.54$ | $-1.23$ |
| Nigeria | $-1.86 \pm 0.44$ | $-2.19 \pm 0.43$ | $-2.50$ |
| Papua New Guinea | $-0.99 \pm 0.69$ | $-1.35 \pm 0.62$ | $-1.20$ |
| Tanzania | $-1.82 \pm 0.45$ | $-2.08 \pm 0.45$ | $-2.47$ |
| Thailand | $-0.96 \pm 0.68$ | $-1.28 \pm 0.61$ | $-1.10$ |
| Vietnam | $-1.49 \pm 0.46$ | $-1.91 \pm 0.47$ | $-1.20$ |

Table 3: Tajima's $D$ across *P. falciparum* populations. Values show mean ± SD of per-gene estimates for synonymous ($D_S$) and nonsynonymous ($D_N$) mutations separately. Genome-wide $D$ calculates across all variable loci in the genome in the population.

| Country | $N_e$ (theta) | $N_e$ (LD) |
|---|---|---|
| *Sub-Saharan Africa (f = 0.3)* | | |
| Nigeria | 69,195 | $10,015 \pm 788$ |
| Tanzania | 83,630 | $9,638 \pm 944$ |
| Cameroon | 83,084 | $9,749 \pm 964$ |
| Kenya | 85,481 | $9,632 \pm 874$ |
| Gambia | 78,306 | $9,325 \pm 779$ |
| Mozambique | 69,985 | $11,134 \pm 783$ |
| Malawi | 86,019 | $9,247 \pm 797$ |
| Mali | 78,224 | $11,104 \pm 973$ |
| Ghana | 78,188 | $10,248 \pm 905$ |
| DRC | 75,363 | $10,822 \pm 938$ |
| *Asia and Oceania (f = 0.7)* | | |
| India | 70,833 | $21,898 \pm 1,893$ |
| Bangladesh | 75,323 | $19,889 \pm 1,842$ |
| Myanmar | 64,373 | $15,375 \pm 1,874$ |
| Thailand | 60,730 | $14,624 \pm 1,862$ |
| Cambodia | 52,926 | $14,856 \pm 1,663$ |
| Laos | 63,665 | $14,851 \pm 1,530$ |
| Vietnam | 49,474 | $12,602 \pm 1,888$ |
| Papua New Guinea | 63,151 | $14,198 \pm 2,035$ |

Table 4: Effective population size estimates for *Plasmodium falciparum* by country. $N_e$-theta estimates are based on nucleotide diversity. $N_e$-LD estimates are corrected for inbreeding using region-specific outcrossing rates: $f = 0.3$ for Sub-Saharan African populations (higher transmission intensity, more outcrossing) and $f = 0.7$ for Asian and Oceanian populations (lower transmission intensity, more selfing), following the formula $E[r^2] = 1/(1 + 2N_e c(1 - f)) + b$. Recombination rate $7.4 \times 10^{-7}$ and mutation rate $1.5 \times 10^{-9}$ per basepair per generation.

| Population | $\mathbb{E}[\pi_N]$ $\times 10^{-4}$ | $\mathbb{E}[\pi_S]$ $\times 10^{-4}$ | % of genes $\pi_S = 0$ | % of genes $\pi_N/\pi_S > 1$ | $\mathbb{E}[\pi_N/\pi_S]$ | $\mathbb{E}[\pi_N]/\mathbb{E}[\pi_S]$ |
|---|---|---|---|---|---|---|
| Mali | 3.31 | 5.70 | 1.3 | 30.3 | 1.54 | 0.58 |
| Ghana | 3.34 | 5.78 | 0.8 | 29.9 | 1.48 | 0.57 |
| DRC | 3.24 | 5.59 | 1.7 | 31.7 | 1.84 | 0.57 |
| Nigeria | 3.36 | 5.69 | 3.8 | 33.3 | 1.58 | 0.58 |
| Tanzania | 3.33 | 5.75 | 3.4 | 32.5 | 1.96 | 0.57 |
| Cameroon | 3.28 | 5.73 | 10.4 | 34.8 | 1.60 | 0.54 |
| Kenya | 3.34 | 5.78 | 3.7 | 34.8 | 2.65 | 0.57 |
| Gambia | 3.28 | 5.66 | 2.1 | 34.6 | 2.25 | 0.58 |
| Mozambique | 3.19 | 5.56 | 2.5 | 33.2 | 1.95 | 0.57 |
| Malawi | 3.40 | 5.86 | 8.2 | 34.3 | 1.53 | 0.54 |
| India | 3.26 | 5.39 | 11.4 | 41.0 | 2.06 | 0.56 |
| Bangladesh | 3.05 | 5.01 | 9.9 | 41.3 | 4.13 | 0.57 |
| Myanmar | 2.60 | 4.31 | 16.5 | 49.3 | 5.87 | 0.53 |
| Thailand | 2.44 | 4.03 | 25.8 | 50.5 | 6.93 | 0.49 |
| Cambodia | 2.35 | 3.69 | 8.1 | 51.8 | 6.87 | 0.59 |
| Laos | 2.56 | 4.19 | 11.5 | 52.1 | 8.63 | 0.56 |
| Vietnam | 2.06 | 3.22 | 7.0 | 52.4 | 7.43 | 0.61 |
| Papua New Guinea | 2.80 | 4.46 | 30.4 | 47.4 | 2.33 | 0.48 |
| Mean ± SD | $2.99 \pm 0.43$ | $5.13 \pm 0.88$ | $9.5 \pm 8.3$ | $39.5 \pm 8.6$ | $3.56 \pm 2.46$ | $0.56 \pm 0.04$ |
| Median | 3.25 | 5.58 | 7.6 | 34.8 | 2.01 | 0.57 |

Table 5: Cross-population summary of $\pi_N$ and $\pi_S$ metrics. $\mathbb{E}[\pi_N/\pi_S]$ is the mean over the per-gene $\pi_N/\pi_S$, and $\mathbb{E}[\pi_N]/\mathbb{E}[\pi_S]$ is the genome-wide mean $\pi_N$ divided by genome-wide $\pi_S$.

| Gene ID | Name | Type | Description |
|---|---|---|---|
| PF3D7_1133400 | AMA1 | Invasion | Apical membrane antigen 1 |
| PF3D7_1035500 | MSP6 | Surface | Merozoite surface protein 6 |
| PF3D7_1035700 | DBLMSP | Surface | Duffy binding-like merozoite surface protein |
| PF3D7_0208900 | P230p | Sexual | 6-cysteine protein P230p |
| PF3D7_0209000 | P230 | Sexual | 6-cysteine protein P230 |
| PF3D7_0830800 | SURFIN8.2 | Variant | Surface-associated interspersed protein 8.2 |
| PF3D7_0402200 | SURFIN4.1 | Variant | Surface-associated interspersed protein 4.1 |
| PF3D7_1301800 | SURFIN13.1 | Variant | Surface-associated interspersed protein 13.1 |
| PF3D7_0937100 | — | Export | Plasmodium exported protein, unknown function |
| PF3D7_0301600 | hyp1 | Export | Plasmodium exported protein (hyp1) |
| PF3D7_1475900 | KELT | Other | KELT protein |
| PF3D7_0320400 | Cap380 | Sexual | Oocyst capsule protein Cap380 |
| PF3D7_0218600 | PNP4 | Other | Patatin-like phospholipase 4 |
| PF3D7_1344800 | — | Metabolism | Aspartate carbamoyltransferase |
| PF3D7_0412300 | — | Metabolism | Phosphopantothenoylcysteine synthetase |
| PF3D7_0211700 | — | Signaling | Tyrosine kinase-like protein, putative |
| PF3D7_1313600 | — | Other | Clu domain-containing protein, putative |

Table 6: Genes with $\pi_N/\pi_S > 1$ across all 18 countries using filtering (variants >0.5% frequency and >5 segregating sites per gene)

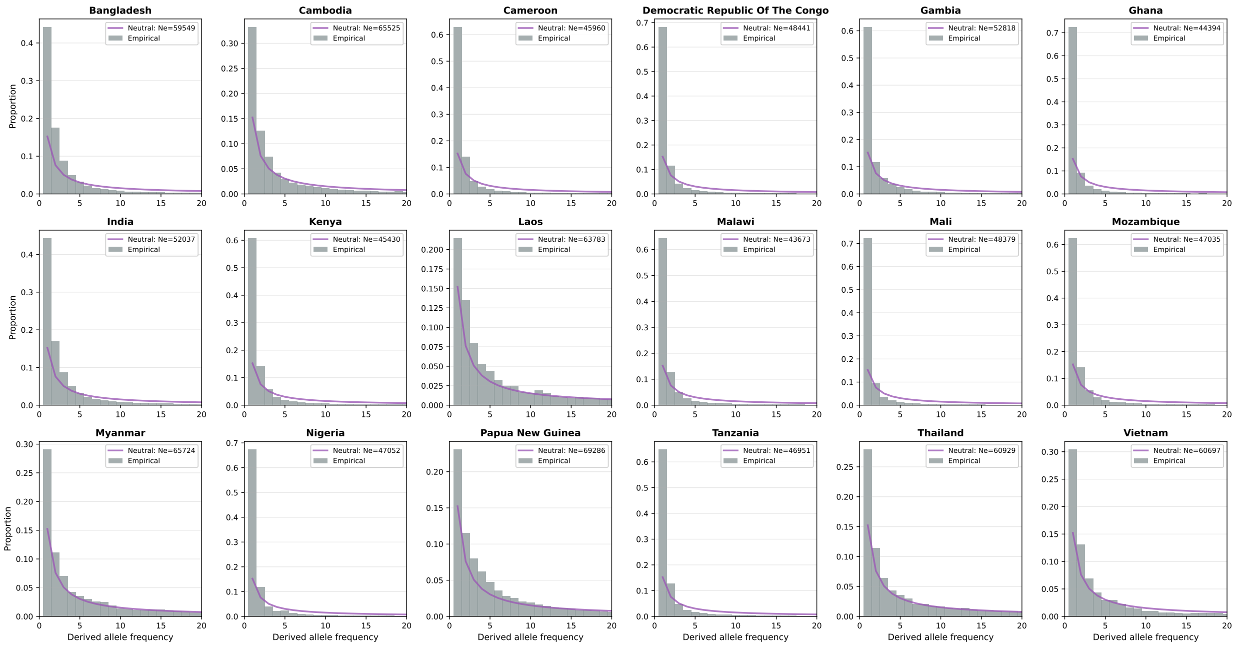


Figure 1: **First 20 bins of the SFS from 18 *P. falciparum* populations compared to neutral Kingman coalescent expectations.** Each panel shows the observed SFS (grey bars) for one population compared to the expected SFS under a constant-size Kingman coalescent (purple line) with effective population size estimated from nucleotide diversity. All populations show pronounced excess of rare variants relative to neutral expectations, with African populations showing the most extreme deviations. The Kingman coalescent systematically underestimates singleton counts while overestimating intermediate-frequency variant counts. Sample size n=200 for all populations.

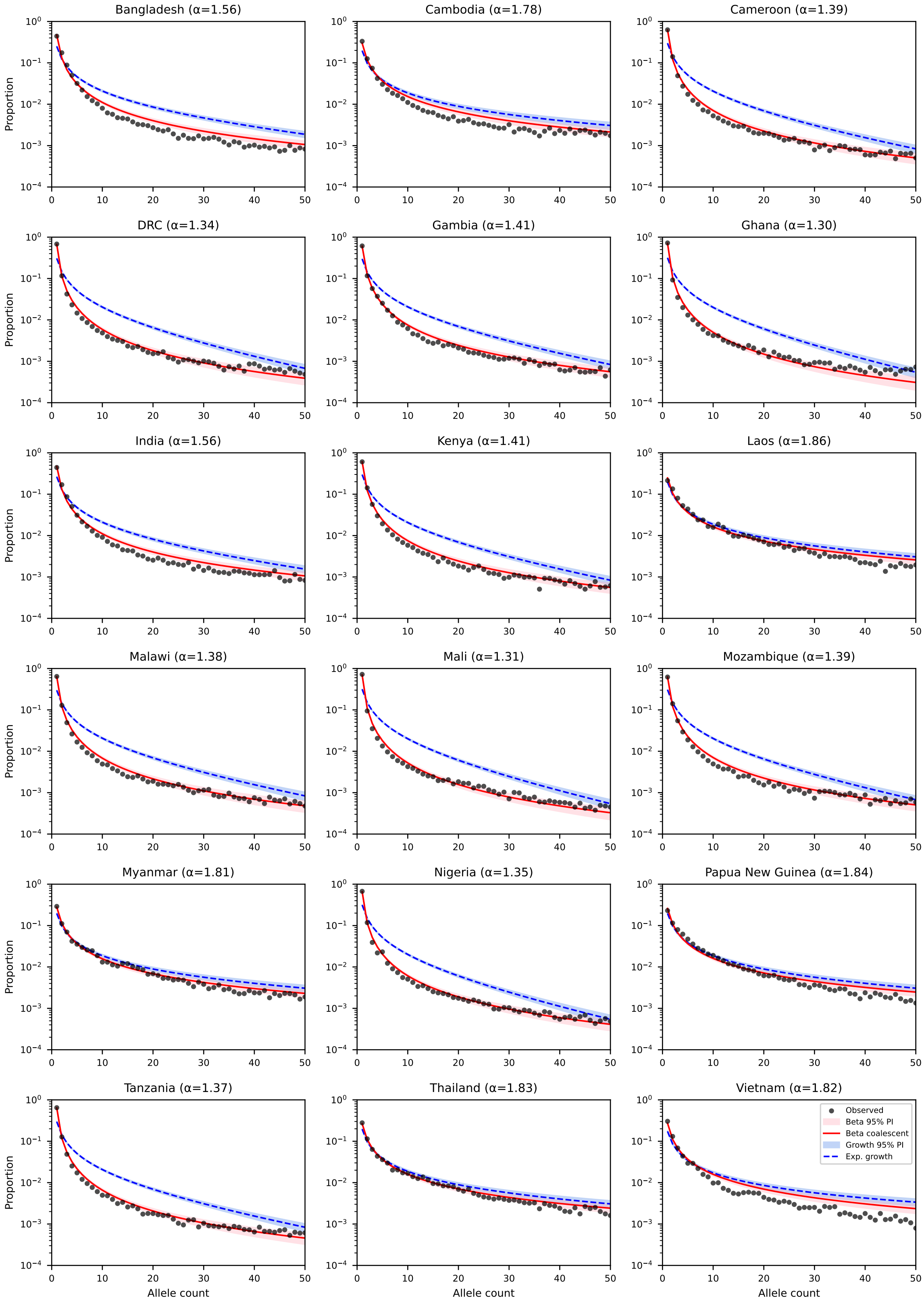


Figure 2: **Comparison of Beta coalescent and exponential growth models for SFS.** Observed SFS (black points) written as a proportion of variants in each bin, with fitted Beta coalescent model (red) and exponential growth model (blue) for each of 18 populations.

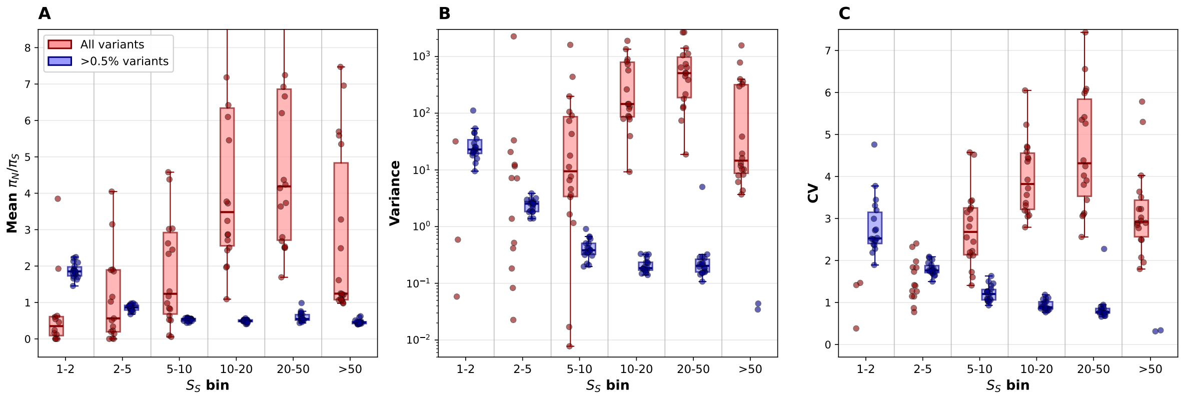


Figure 3: The distribution of gene-level $\pi_N/\pi_S$ ratios stratified by number of synonymous segregating sites with and without frequency filtering. **(A)** Mean $\pi_N/\pi_S$ by segregating site bin for all variants (pink) versus variants >0.5% frequency (blue). Without filtering, mean ratios remain elevated regardless of segregating site count, contradicting theoretical predictions. After frequency filtering, ratios decline with increasing segregating sites as predicted by sampling variance theory. **(B)** Variance of $\pi_N/\pi_S$ by segregating site bin, showing dramatic reduction after filtering, particularly for low-diversity genes. **(C)** Coefficient of variation ($\mathbb{CV}$) by segregating site bin. Points represent individual countries.

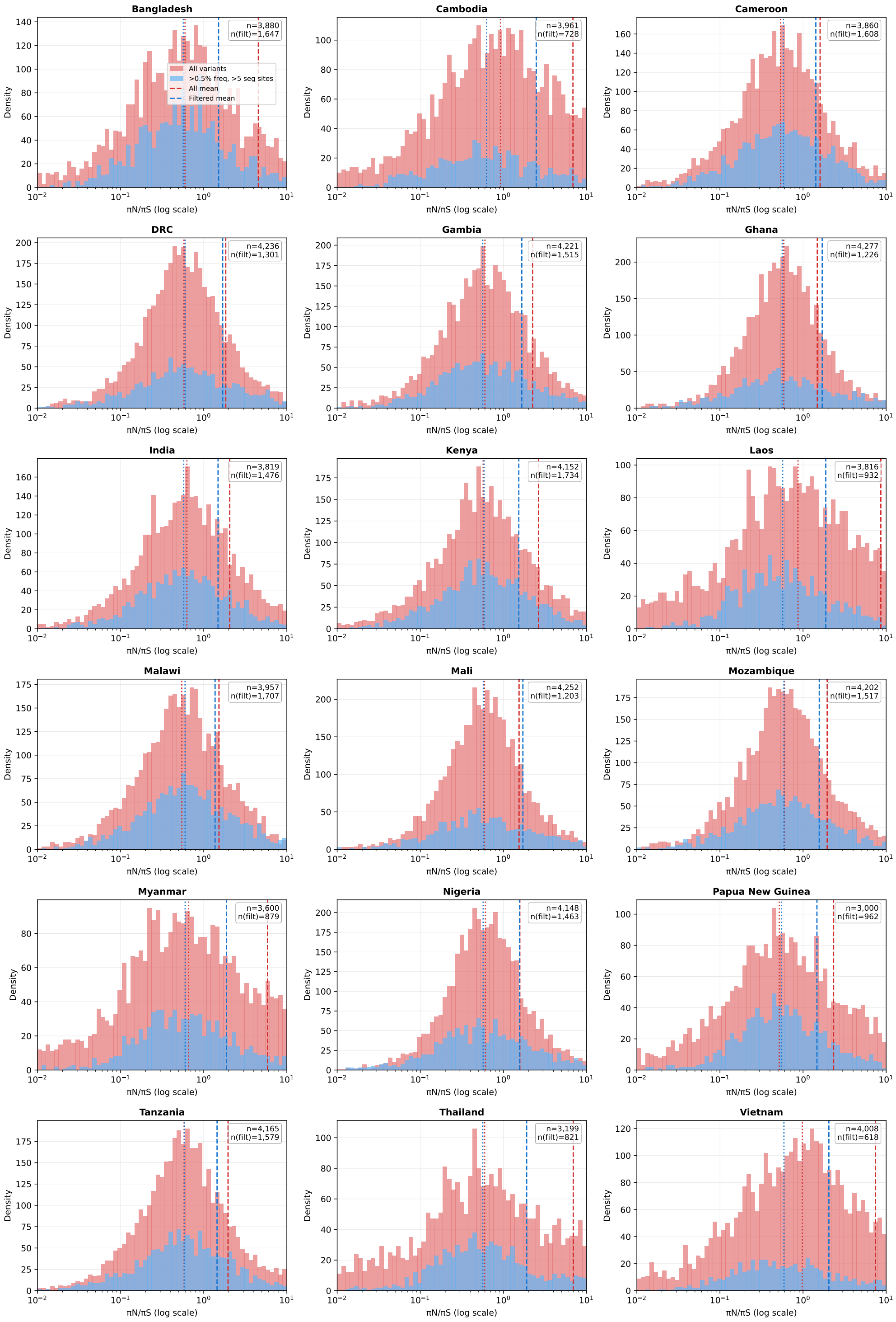


Figure 4: Distribution of per-gene $\pi_N/\pi_S$ ratios across 18 countries. Red histograms show ratios calculated from all segregating variants regardless of frequency (n = 3,000-4,277 genes per country). Blue histograms show ratios calculated after filtering for variants with >0.5% minor allele frequency and genes with >5 segregating sites (n = 618-1,734 genes per country). Dashed vertical lines indicate mean values for each distribution; dotted lines indicate medians. Sample sizes for total genes (n) and filtered genes are shown in each panel. Unfiltered datasets show inflated $\pi_N/\pi_S$ ratios with high variance and right skew. After filtering, all countries converge to similar, biologically plausible $\pi_N/\pi_S$ ratios centered around 1.4-1.9, with substantially reduced variance.

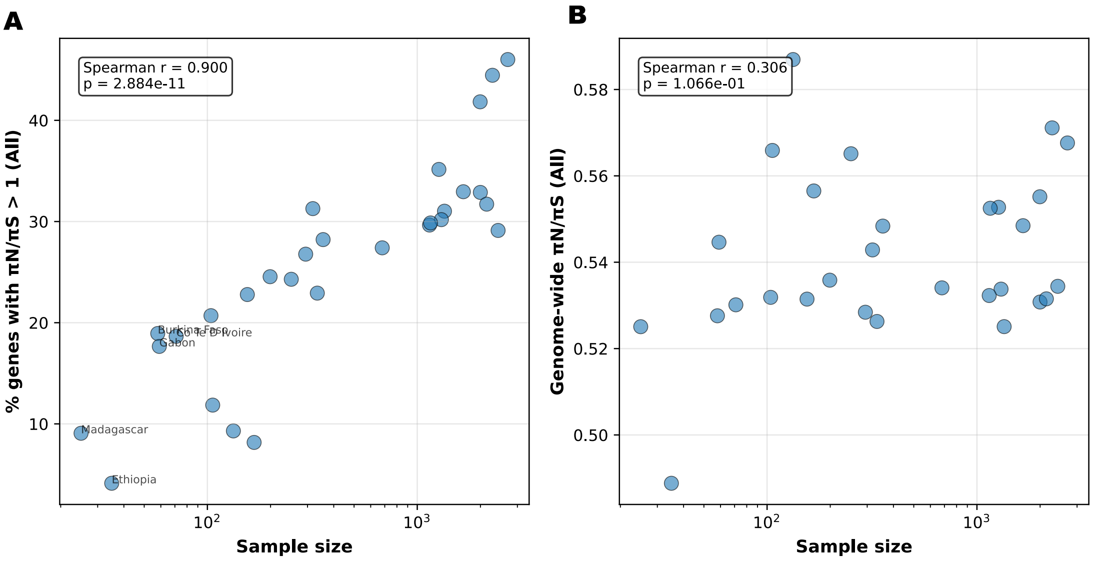


Figure 5: Sample size is correlated with the proportion of genes with $\pi_N/\pi_S > 1$, but not with genome-wide $\mathbb{E}[\pi_N]/\mathbb{E}[\pi_S]$. Data from 29 countries from MalariaGEN $Pf8$. Further analyses focused on the 18 countries with at least 200 samples.
